# MHC class I and II genes in Serpentes

**DOI:** 10.1101/2020.06.12.133363

**Authors:** D.N. Olivieri, S. Mirete-Bachiller, F. Gambón-Deza

**Affiliations:** Department of Computer Science, University of Vigo, Ourense 32004, Spain; Department of Immunology, Hospital of Meixoiero, Vigo, SPAIN

**Keywords:** Major Histocompatibility Complex, Serpentes, Immunity Reptiles

## Abstract

Genes of the major class I and II histocompatibility complex have been extensively studied in mammals. Studies of these antigens in reptiles are very scarce. Here we describe the characteristics of these genes in the suborder Serpentes. We identified the presence of a much larger number of molecules of class I and beta chains of class II than found in mammals. Snakes only have one gene for the class II alpha chain. In these species, class I genes can be classified into two types. Approximately half of the genes lack 10 amino acids in the *α*1 domain, producing a structural alteration in the interaction region with the T lymphocyte receptor. In the genome of *Thamnophis elegans*, two haplotypes of an individual were studied revealing a different number and location of class I genes between these haplotypes. The results indicate that in these species, the diversity in the MHC is generated by the presence or absence of genes, independent of the presence of alleles.

## 1. Introduction

The major histocompatibility complex (MHC) was described as the genome region that encodes proteins responsible for organ transplant rejection (Zinkernagel & Doherty, 1997). Over time, these proteins were described. They correspond mainly to integral membrane proteins with amino acid sequences belonging to the immunoglobulin family. Studies about their function revealed that they possess a great amount of allelic polymorphism and that they play an important role in antigen presentation to T lymphocytes (Neefjes et al., 2011). Two classes were described. Class I (MHC-I) is made up of a main polypeptide chain that is non-covalently linked to *β*2-microglobulin (Collins et al., 1994). The first two protein domains arrange to form a cleft that can harbor peptide fragments to be presented to CD8+ T lymphocytes. This class mainly presents cytosolic peptide fragments produced by protein alterations (Engelhard, 1994). Class II (MHC-II) is made up of two polypeptide chains, such that the outermost domains also arrange to form a cleft structure similar to that of the class I antigens, but able to bind slightly larger peptide fragments. These molecules present foreign antigens that have been endocytosed and degraded by cells. These fragments are presented to CD4+ lymphocytes for triggering immune responses (Doyle & Strominger, 1987).

These molecules appeared as such with the rise of jawed vertebrates and occurred in conjunction with genes for immunoglobulins and T-cell receptor genes (Okamura et al., 1997). These antigens have been extensively studied in mammals, revealing that there are several genes for class I molecules and all are similar to each other. A group of these antigens have non-specific tissue expression, and often large allelic variation; these are referred to as classical MHC antigens. The other group of antigens are expressed in specific tissues and do not display allelic variation; such antigens are called non-classical MHC antigens (Watkins et al., 1991). For the conformation of class II antigens, one gene for the alpha chain and one for the beta chain are required. In mammals, there are various gene pairings for alpha and beta. Sometimes the alpha chain gene is accompanied by several beta chain genes (Takahashi et al., 2000). Class II molecules are exclusively expressed in specialized cells for antigen presentation (Ting & Trowsdale, 2002). Their participation in the defense against infection has been clearly demonstrated. Currently, the field is much broader since it is believed that the immune response also participates in anti-tumor defense and the elimination of senescent cells, functions that may have preceded that of the defense against infection.

Very little data exists on MHC in reptiles (Elbers & Taylor, 2016). There are some descriptive publications on the sequencing of mRNA and the deduced proteins in Iguanas (Glaberman et al., 2008, 2009) and Sphenodontia (Miller et al., 2006). More recently, these molecules have also been described in Cocodrylia (Jaratlerdsiri et al., 2014), however there is no comprehensive description of the number of genes and their location in the genome in these species.

In this article, we present the MHC genes obtained from genomes of the suborder Serpentes using a bioinformatics approach. By comparing this data with other species within and outside this reptile suborder, structural differences have been observed. More broadly, the study of the evolution of these MHC molecules may reveal interesting clues about their specific function and the origins of the immune system.

## 2. Methods

### 2.1. Datasets

The genome sequences were obtained from the NCBI and the geneArk project (https://vgp.github.io/genomeark-curated-assembly/). The RefSeq annotations of these genomes were obtained from the NCBI public repository. The software analysis code and sequences obtained used in this work are available at http://vgenerepertoire.org/ and deposited on the public data repository Dryad.

### 2.2 Software

While there are six viable MHC-I genes in *Homo sapiens*, the number of MHC-I genes varies considerably across other mammal species. Nonetheless, the MHC-I exon architecture is nearly universal across all mammal species (Birch et al., 2006). A diagram of the *Homo sapiens* MHC-I exon structure and corresponding protein domains was described (Lefranc et al., 2005) (see also sequence repositories of the by the IMGT) and the a large collection of MHC-I alleles at the IPD-MHC database (Robinson et al., 2011). The relevant genomic signals (i.e., the exons, introns, 5′-URT and 3′-URT) can be identified to a high degree of accuracy without resorting to general gene finding software tools. Therefore, viable exons (i.e., those exons that could form a functionally expressed MHC-I molecule) can be accurately determined using homology similarity with a supervised machine learning classifier.

The `MHCfinder` program extracts exons EX2, EX3, and EX4 (from the IMGT nomenclature for Exon-2, Exon-3 and Exon-4) that encode the *α*1, *α*2 and *α*3 domains. Although other exons are required to constitute the full MHC-I gene, these three exons are of particular interest for characterizing and comparing MHC-I genes within and amongst species. Thus, `MHCfinder` ignores the peptide leader (L), as well as all peptides corresponding to the transmembrane and cytoplasmic (*T_m_*). While the exon/intron structure of MHC Class I is thought to be universal across jawed vertebrates, the specific intron spacing between the exon sequences EX2, EX3, and EX4 varies considerably. As such, the algorithm only imposes a simple structural requirement that EX2, EX3 and EX4 are found in a tandem arrangement along the DNA sequence, but places no hard restrictions on the intron separation.

Figure 1 summarizes the principal steps of the `MHCfinder` algorithm. This program was implemented as a multi-threaded application in the python programming language, with the biopython library (Cock et al., 2009) for low-level sequence analysis, and both the scikits library (Pedregosa et al., 2011) and TensorFlow framework (Abadi et al., 2016) for machine learning tasks. First, a Tblastn query from a consensus protein sequences from known MHC-I and MHC-II exons is made against all available genome datasets. The search result is a listing of candidate WGS contigs likely to contain valid exons, together with the position of the matching nucleotide sequence and similarity scores; this listing is referred to as a *hit table*. The algorithm processes each line of the *hit table*, analyzing a nucleotide region larger on both ends than the nucleotide positions in the hit. Then a more precise algorithm is used to enumerate all the potential exon reading frames defined by the positions of the start/stop motifs, `AG` and `GT`, respectively.

**Figure 1:**
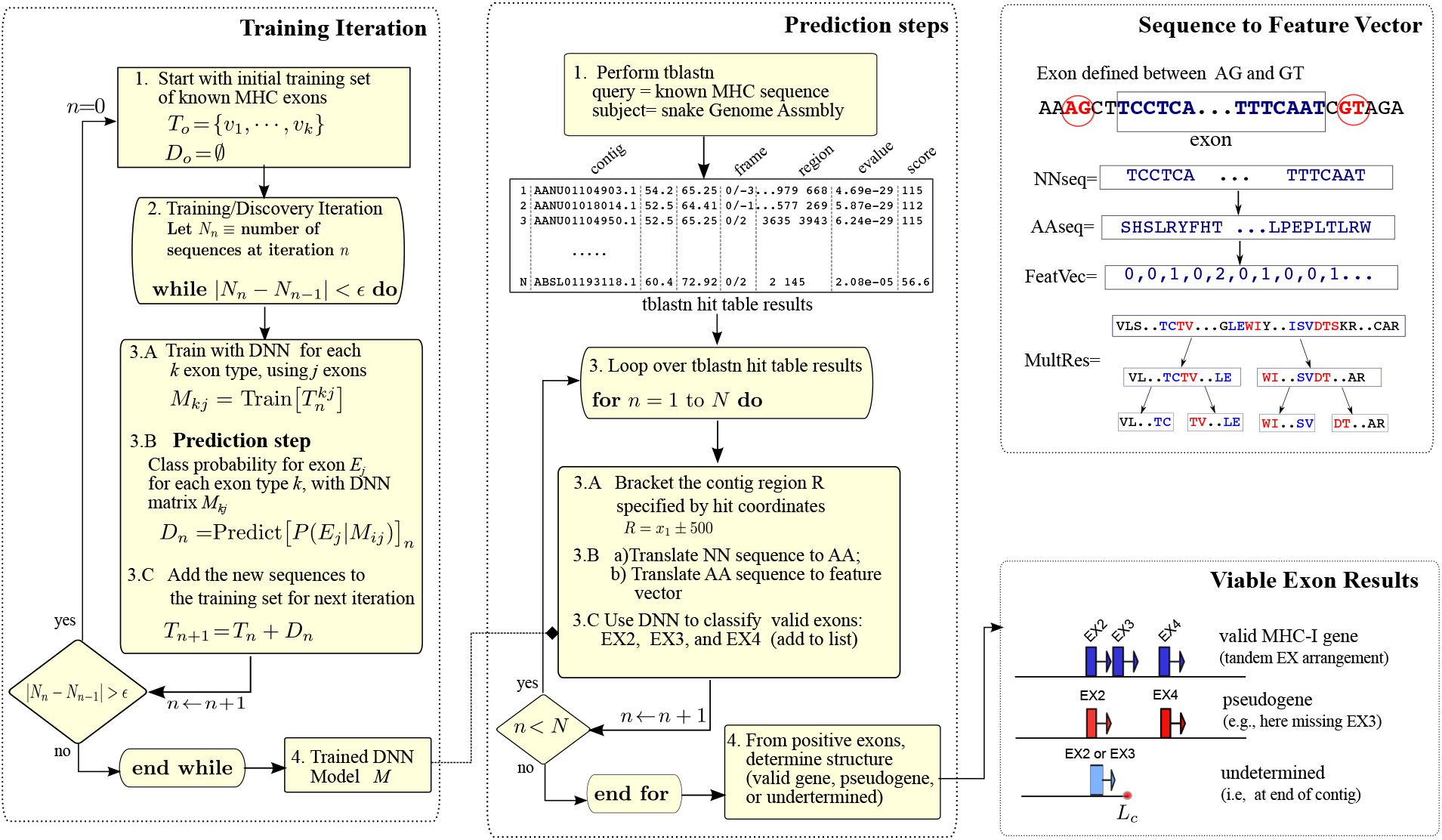
The steps in the MHC training and prediction algorithms. The training phase starts with known MHC exons from other jawed vertebrates, and iteratively includes newly discovered sequences into the training set. The result is a trained Deep Neural Network (DNN) model, *M*. In the first part of the prediction phase, locations of potential MHC-I and MHC-II exons within contigs are found from a Tblastn query. These queries are only used for bracketing a finer search. In this way, all possible reading frames of candidate exon are obtained from these regions and evaluated with the DNN model *M*, to determine if they are valid and which has the largest score. This process also chooses the best candidate from overlapping exons. As a result of this process, valid sequences are classified into their respective exon class. Tandem exons EX2-EX3-EX4 are identified and used to construct viable MHC-I or MHC-II genes.

Once candidate exons are identified, they are translated into amino acid sequences by checking all valid reading frames. Those sequences containing stop-codons in the reading frame are discarded, while valid exons are saved and converted into numerical feature vectors (i.e., a unique array of numbers, that characterizes the string of amino acids). Two separate coding schemes were used for independent checks: a multi-resolution method (seen in Figure 1 and a linear coding. Both methods are based upon the individual and co-occurring amino acid frequency tuples (ie. tallying the sequence and frequency of each AA and pairs of AA). In our studies, these custom encoding schemes proved more effective and accurate for homology detection than published methods based upon positional physicochemical properties of each AA within the sequence.

From the feature vector representation, a machine learning procedure with a Deep Neural Network (DNN) using TensorFlow was used to classify the sequence into one of the exon types: EX2, EX3, and EX4. Multi-class supervised learning is used to train the DNN model with these exon classes from annotated exons and a random background sequences. For completeness, we compared and confirmed the results from the DNN classifier with a random forest method, as well as hierarchical clustering. The classification is carried out for each exon type with a background/signal ratio of 3:1. The prediction precision was improved by multiple training/prediction iterations; positively identified sequences are included in the training set for subsequent training/predictions (see (Olivieri & Gambón-Deza, 2019) for details of the method). This process is referred to as iterative supervised learning, and is a common machine learning technique whereby new information is continually accrued to the knowledge base for improving prediction accuracy.

In our gene finding algorithm, `MHCfinder`, a probable functionally expressed MHC-I gene that must contain a tandem arrangement of the three viable exons EX2-EX3-EX4 without stop codons and with canonical intron distances. Nonetheless, viable exons, which are homologous to either MHC-I constituent exons, exist in the genome that do not form tandem arrangements (i.e., they are isolated or an exon is missing), and thus, do not express MHC molecules.

The same procedure was used to identify MHC-II antigens. For this, `MHCfinder` was trained with annotated sequences of the *α*1, *α*2 (the alpha chain of class D, DM and DO) domains and *β*1, *β*2 (the beta chain of class D, DM and DO) domains. The machine learning procedure achieves sequence identification results with an accuracy greater than 99%.

#### Tree construction

To study the phylogenetic relationships from the MHC-I exons, we constructed a large phylogenetic tree by aligning sequences with ClustalO and then used phyML with the LG matrix (part of the Fasttree software (Price et al., 2010)). In all cases, 500 bootstrapped samples were made.

#### 3D model

Three-dimensional structure models of snake MHC-I antigens were obtained from amino acid sequences obtained from predictions with the Swiss-Model ExPASy (found at https://swissmodel.expasy.org/)

## 3. Results

To search for MHC class I and II genes, we used a double strategy. We started from the existing annotation in the NCBI of reptile genomes obtained from the RefSeq repository. We selected those sequences having the word histocompatibility in their descriptions. Additionally, these sequences were used together with sequences from other vertebrates to carry out machine learning that would recognize the main exons of these molecules. The combination of both methods gave us concordant results for the study.

### Thamnophis elegans

Although many of the genome sequencing projects for reptiles are in an advanced state, the MHC genes appear in several scaffolds in most of these genomes and not yet assigned to chromosomes. In the genomes for the snake *Thamnophis elegans*, most MHC genes are on one chromosome. All the sequences were found in two segments: all the genes for MHC-I and some for MHC-II are found in the Super_Sacaffold_1 (nomenclatura de VGP, NCBI NC 45542.1), while those for the beta chain of MHC-II together with the genes for the DM were found in scaffold 59 arrow ctg1 (NCBI NW 022473899.1).

The RefSeq file for *Thamnophis elegans* is available at the NCBI. It contains 56 sequences that are annotated as major histocompatibility complex class I-related. Some of the sequences are isomorphic to the same gene, so the number of genes identified is somewhat less. With machine learning we identified 201 possible exons for exons EX2, EX3 and EX4 of the genes for MHC-I (for domains *α*1,*α*2 and *α*3). A schematic representation is shown in figure 2. As described in the methods section, tandem exons EX2-EX3-EX4 are identified for deducing and building viable MHC-I. With this method, we identified 65 MHC-I genes. Most of the sequences obtained from the RefSeq are coincident with the sequences obtained with the `MHCfinder`. An explanation of why more sequences are found with the computer program is expressed later in the discussion.

**Figure 2:**
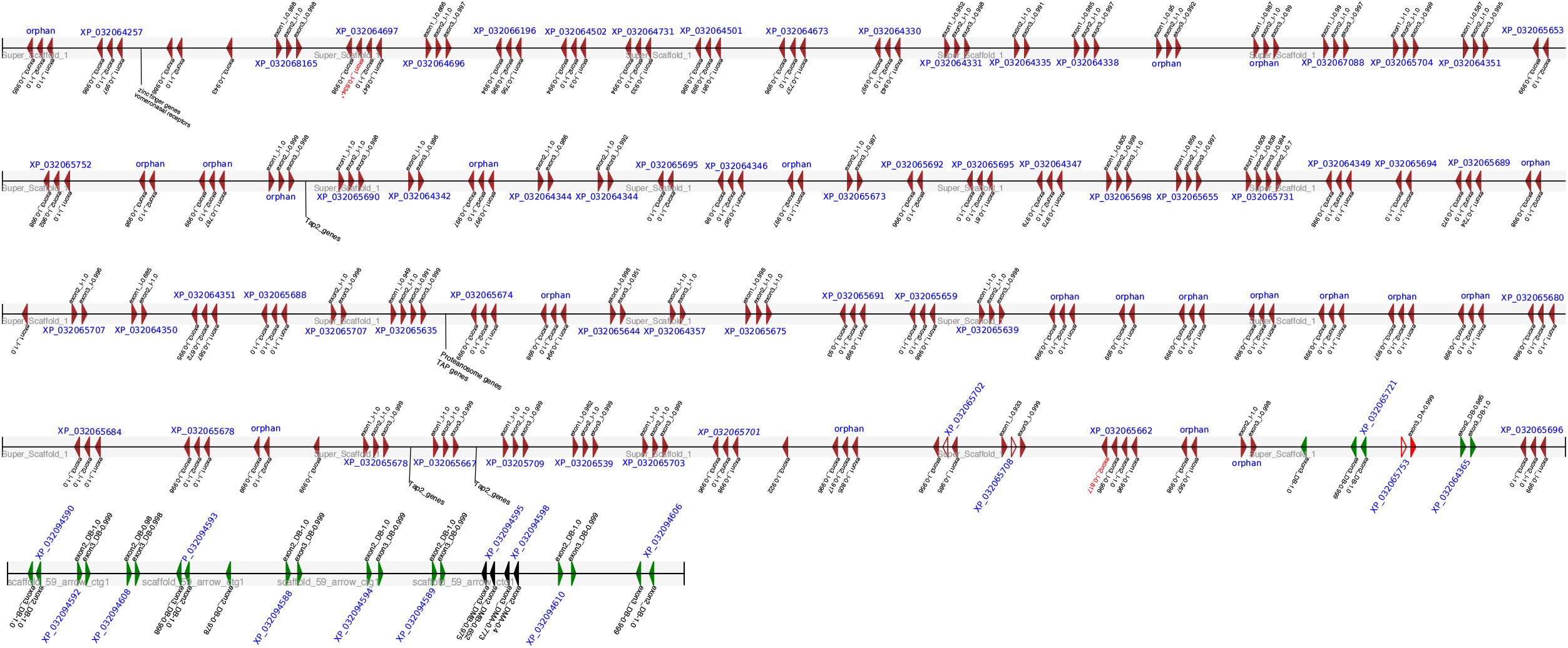
Schematic diagram of the main exons of the MHC antigens found with the MHCfinder program. The different coding exons are shown corresponding to the different domains: (brown) the exons coding for the MHC-I *α*1, *α*2 and *α*3 domains, (red) the coding exons for the MHC-II *α*1 and *α*2 domains, (green) the DB exons *β*1 and *β*2, and (black) the DM exons for *α*1, *α*2/*β*1, *β*2. For each exon, the prediction probability is indicated. For a set of contiguous exons, the NCBI annotation of the RefSeq protein sequence (i.e., identifier starting with XP) is provided. Those groups not found by the NCBI and notated in RefSeq are indicated as NF.

The MHC-II antigens have two chains. With `MHCfinder` we only found one gene for the alpha chain and 11 genes for the beta chain. In this case, the results from RefSeq and from our machine learning procedure coincided. In addition, one gene was found for the alpha chain and another for the beta of the invariant DM genes. No genes were found for DO. The beta-chain genes have similar sequences and different classes are not distinguished as in mammals (DQ,DP, DR).

We studied whether the presence of a high number of MHC-I genes and MHC-II beta chains is particular to this species or can be generalized to all snake species. With the `MHCfinder`, we search for the presence of the seven major exons in the genomes of other snakes available on the NCBI public repository (table 1). The MHC region in most genomes is only partially assembled and contains gaps, so the results we found are approximate. A large number of MHC-I exons are detected in all of the species and there is a single gene for the MHC-II alpha chain and multiple genes for the beta chain can be deduced. From this, it can be concluded that all snakes share an organization similar to that described in *Thamnophis elegans*.

**Table 1:** Number of major exons for MHC-I and MHC-II in the suborder Serpentes

| SERPENTES | ex1_I | ex2_I | ex3_I | ex2_DA | ex3_DA | ex2_DB | ex3_DB |
| --- | --- | --- | --- | --- | --- | --- | --- |
| *Thamnophis elegans* | 69 | 83 | 87 | 1 | 2 | 13 | 12 |
| *Thermophis baileyi* | 23 | 28 | 30 | 1 | 3 | 18 | 17 |
| *Pseudonaja textilis* | 34 | 28 | 35 | 1 | 1 | 22 | 8 |
| *P. mucrosquamatus* | 72 | 81 | 83 | 1 | 2 | 86 | 83 |
| *Notechis scutatus* | 39 | 38 | 35 | 1 | 2 | 26 | 25 |
| *Naja naja* | 52 | 62 | 62 | 1 | 2 | 22 | 23 |
| *Crotalus viridis* | 14 | 15 | 20 | - | - | 3 | 8 |
| *Laticauda colubrina* | 13 | 21 | 19 | - | 1 | 5 | 10 |

The sequences of the MHC-I antigens generate two very differentiated evolutionary clades (figure 3). In one clade, a significant alteration in sequences from the *α*1 domain exists, in which a 10 amino acids long peptide has been deleted. In the other clade, this same amino acid peptide region is highly conserved in all the sequences. These results suggest a separate functions between the two clades.

**Figure 3:**
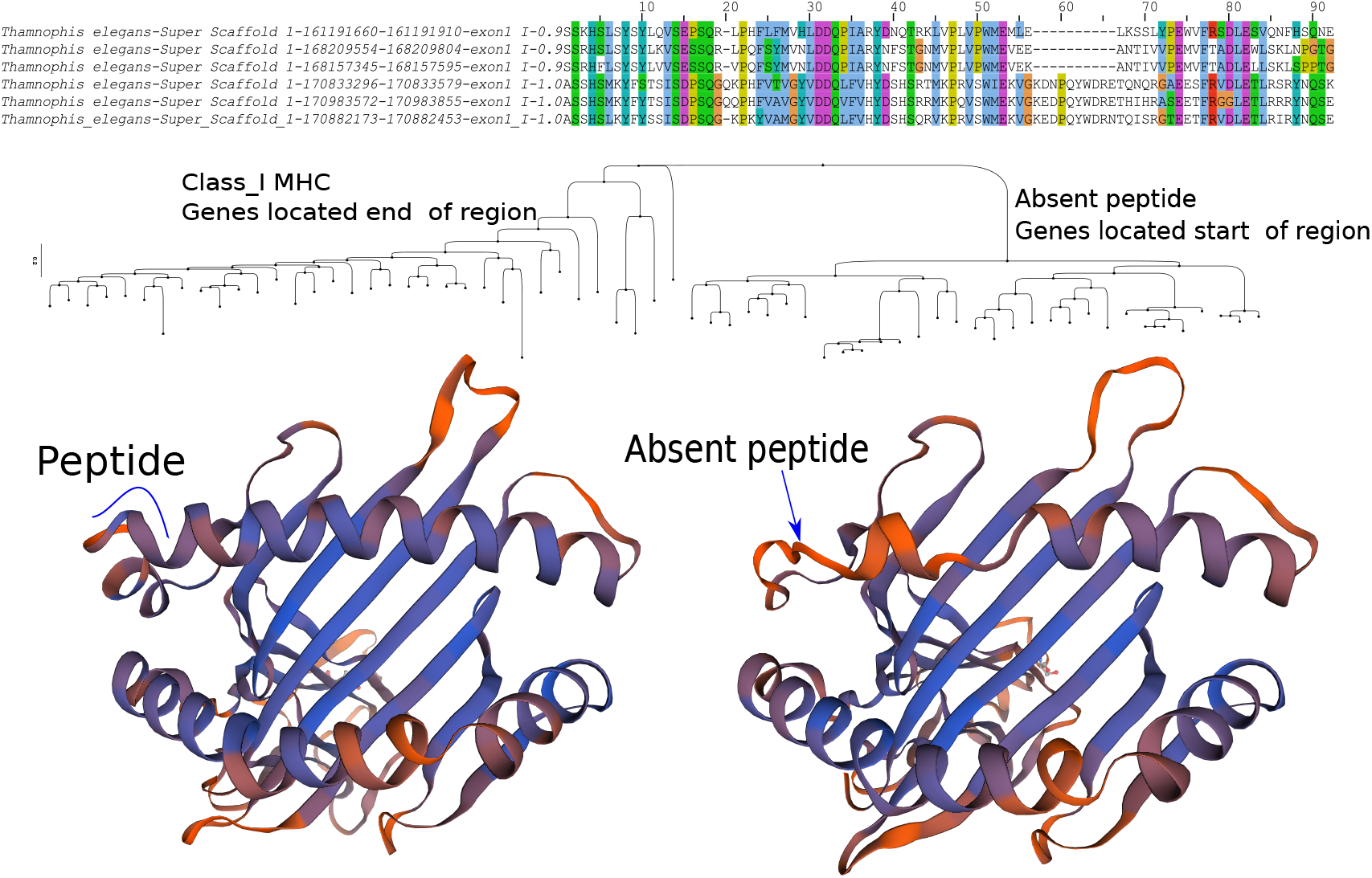
Phylogenetic trees were made with the deduced amino acid sequences of the genes obtained in this study. Two clades are evident. There is a structural difference in the sequences of each clade due to the absence of a 10 amino acid long peptide in the *α*-1 domain. Alignment of three *α*1 domain sequences from each group (having peptide and the absent peptide group) are shown at the top of the figure. The three-dimensional representations were obtained with a sequence of each clade with Swiss-Model ExPASy predictions.

To explore the structural implications of this peptide variation, we obtained 3D structure predictions of chains from each clade with Swiss-Model ExPASy program (https://swissmodel.expasy.org/). Figure 3 shows the results of representative sequences from each clade. The 10 amino acid long peptide under study is located at the start of the first alpha helix. Its absence radically alters the canonical alpha helical structure. Considering that this zone is a docking site for the CDRs of the TCR chains, it may be indicative of a fundamental functional alteration.

The absence of this peptide at the start of the alpha helix is also found in other snakes (*Naja naja*, *Notechis scutatus*, *Pseudonaja textilis* and *Crotalus viridis*). Figure 4 shows the tree generated with the sequences for the *α*1 domain. The clade with the absent peptide sequence contains sequences of the five species. This indicates that the generation of these molecules predates the divergence of the snake species, suggesting that all members of this suborder must possess this class of MHC-I *α*1 domain.

**Figure 4:**
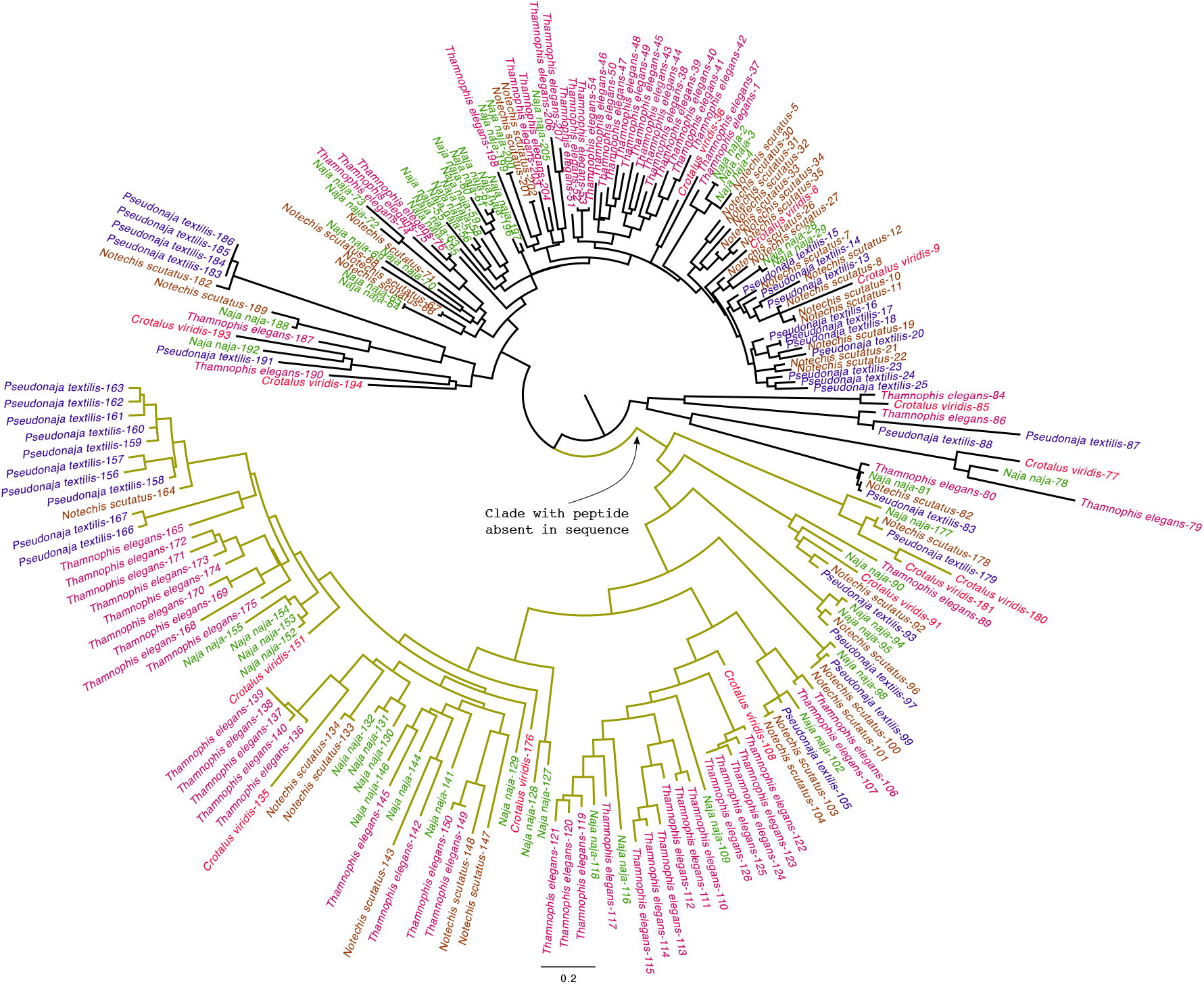
The phylogenetic tree made with the deduced amino acid sequences of the exons coding for the domain *α*1 of the MHC class I molecules from 4 snake species. Each species is colored differently for clarity. The clade with the altered MHC-I *α*1 domain (with absent peptide) referred to in the text is indicated.

In the genome of *Thamnophis elegans*, both haplotypes have been obtained from one individual. By comparing the two haplotypes, we found that there is no agreement between the genes. In the alternative haplotype, we selected the longest contigs and aligned them with the primary haplotype. A dotplot of sequences for several genes of MHC-I is shown in Figure 5(left). The presence of overlapping duplications and sequence segments between haplotypes is evident. Furthermore, in some regions there are genes that are found in one haplotype and not in the other. Figure 5(right) shows the dotplot constructed with MHC-II genes from a single contig. In this case, the sequences are more similar, and have the same total number of genes. These results seem to suggest that the mechanism for generating individual diversity of MHC-I genes in snakes is different from that of mammals.

**Figure 5:**
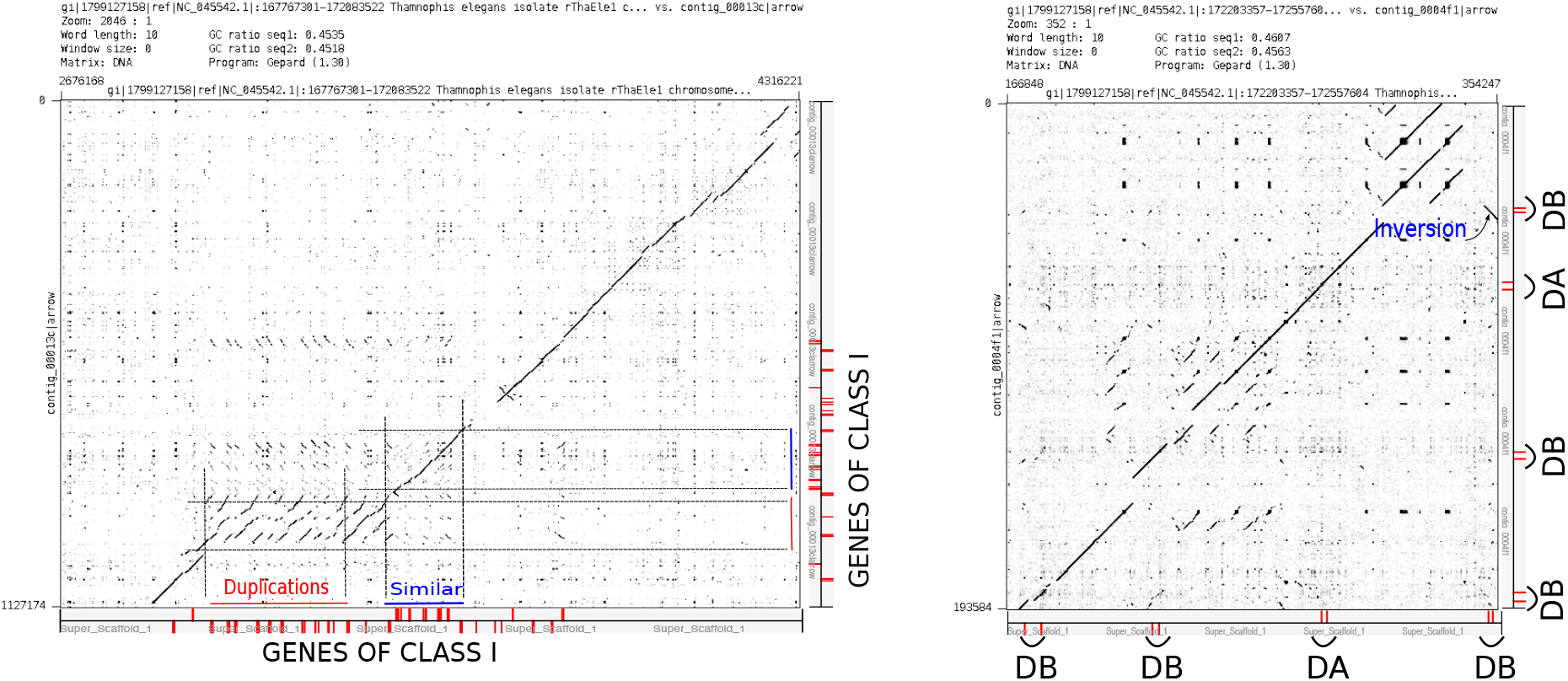
(Left) MHC-I dotplot of the contig 00013c from the alternative genome versus the chromosome region located at the end of the locus. The location of the coding exons for the three main MHC-I domains are indicated in red on the axis of the graph. Areas with low similarity (red) and exact matches (blue) are indicated with boxes. (Right) MHC-II dotplot of the contig 0004fi from the alternative genome versus the chromosome region located at the end of the locus. The coding exons for MHC-II antigens are indicated on the axes of the graph.

## 4. Discussion

There are few studies that describe the major histocompatibility complex in reptiles. Here, we present data focused on class I and II (MHC-I and MHC-II) antigens in Snakes. The large number of genes for the MHC-I antigens is notable and highly conspicuous. The data from *Thamnophis elegans* correspond to the largest number of MHC-I genes found so far in a vertebrate species. The 75 genes described are grouped into two clades. While one of the clades corresponds to proteins very similar to those described in mammals, a second clade consists of sequences that are missing a 10 amino acids long peptide region that is within the interaction site of TCR and the presented antigen peptide. This structural difference may be related to the surprisingly low number of V*β* regions present in these species. Previous publications have already demonstrated the absence of TCR *γ/δ* in snakes and the high number of V *α* genes and only 1 to 5 V*β* genes (Olivieri et al., 2014). Peptide and TCR docking studies may clarify these questions in the future.

The clade consisting of sequences that lack this 10 amino acid long peptide from the canonical *α*1 domain was generated prior to snake divergence and was apparently positively selected and maintained. In all the snakes in this study, we found these altered MHC-I *α*1 domain sequences. In other reptile species, such sequences were not found. The presence of two clearly differentiated MHC-I types, having alterations in the interaction region with TCR, could be suggestive of different functions for these molecules that are yet to be discovered.

In humans and mice, a distinction has been made between classical and non-classical MHC. At present, sufficient data doesn’t exist to classify snake genes into such categories if they exist. The present study was performed with data from a specific animal, so we cannot define the allelic variation. Nonetheless, the results presented here regarding what was found in the haplotypes of this individual are surprising. In particular, that no exact sequence matches are found indicates that there are different numbers of genes between haplotypes. Having both a large number of MHC-I genes and different number of such genes between haplotypes may be the functionally equivalent to possessing a large allelic variation, as seen in mammals.

These haplotype difference results may also explain the technical problems experienced when sequencing this region. In the genomes deposited in the NCBI, this region is usually divided into numerous contigs, indicative of the difficulty of the sequence assembly process. The presence of non-coincident haplotypes in the sequences may be at the root of this problem. The same is true for obtaining the RefSeq assemblies. The NCBI identify genes from alignments of RNAseq against the genome. Considering that RNAseq are often from samples taken from separate individuals, it would explain the differences found between what is deduced from the genome (high number of genes) and the number of genes deduced from the RNAseq.

There are publications about the allelic variation of MHC in some reptile species (Elbers & Taylor, 2016). These studies suffer from the fact that they were carried out without knowing the number of genes present in the genome. The studies deduced the presence of high polymorphism in these molecules, leaving in doubt whether the polymorphism was due to the presence of multiple genes or to allelic variation. In this work, we conclude that the diversity of MHC antigens in snakes is produced by the large number of genes and variations in the number of these between haplotypes.

In MHC class II genes, the existence of a single gene for the alpha chain and the presence of multiple genes for the beta chain is notable. The results indicate that the generation of population diversity must depend on the beta chain genes. The analysis of haplotypes showed less variability of class II than that of class I, therefore it is inconclusive whether these molecules have the same characteristics for diversity generation.

In conclusion, we describe the presence of a high number of MHC class I genes and class II beta chain genes. The high number of genes and the differences between haplotypes is the source of individual diversity of these molecules in Serpentes. Also, the presence of two types of class I antigens indicate the existence of differentiated functionalities for these molecules. It is therefore expected that these results should have implications on the V gene repertoire of the TCR chains in these species.

